# Anterolateral entorhinal-hippocampal imbalance in older adults disrupts object pattern separation

**DOI:** 10.1101/162925

**Authors:** Zachariah M. Reagh, Jessica A. Noche, Nicholas J. Tustison, Derek Delisle, Elizabeth A. Murray, Michael A. Yassa

**Affiliations:** Department of Neurobiology and Behavior, Center for the Neurobiology of Learning and Memory, UC Institute for Memory Impairments and Neurological Disorders, University of California, Irvine; Department of Radiology and Medical Imaging, University of Virginia, Charlottesville,VA

## Abstract

The entorhinal cortex (EC) is among the earliest brain areas to deteriorate in Alzheimer’s disease (AD). However, the extent to which functional properties of the EC are altered in the aging brain, even in the absence of clinical symptoms, is not understood. Recent human fMRI studies have identified a functional dissociation within the EC, similar to what is found in rodents. Here, we used high-resolution fMRI to identify a specific hypoactivity in the anterolateral EC (alEC) commensurate with major behavioral deficits on an object pattern separation task in asymptomatic older adults. Only subtle deficits were found in a comparable spatial condition, with no associated differences in posteromedial EC between young and older adults. We additionally link this condition to previously reported dentate/CA3 hyperactivity, both of which were associated with object mnemonic discrimination impairment. These results provide novel evidence of alEC-dentate/CA3 circuit dysfunction in cognitively normal aged humans.

## Introduction

The entorhinal cortex (EC) is among the first and most profoundly affected regions in age-related pathologies, such as Alzheimer’s disease (AD)^1-4^. In particular, the “transentorhinal region” comprised of the perirhinal cortex (PRC) and adjacent EC, undergoes structural and functional alterations ^5-9^, even in preclinical human subject^10^.

The EC is a key region of the brain’s medial temporal lobes (MTL), and mediates much of the communication between the hippocampus and the rest of the neocortex. Input from the EC to the dentate gyrus (DG) hippocampus is thought to play a role in pattern separation, a process that involves storage of similar experiences using unique memory traces^11^, and which is disrupted in the aging brain^5,12-15^. Electrophysiological^16,17^ and lesion^18^ studies in rodents have shown a functional dissociation between the lateral and medial EC, with the former comprising a “content” pathway representing local sensory cues required for object processing, and the latter comprising a “context” pathway representing configurational aspects of experience such as space^19-21^. This dissociation has been corroborated in human fMRI studies with task-free time series correlations^22,23^ and task-driven engagement^24,25^. Importantly, in the human brain, EC seems to more closely follow an anterolateral versus posteromedial functional division (alEC; pmEC)^22,23^, which is consistent with anatomical properties of the nonhuman primate^26^.

As the “transentorhinal region” is more characteristic of alEC than pmEC, the hypothesis arises that alEC should be more affected than pmEC as memory declines with aging. This is in line with findings by Khan and colleagues^10^, who reported metabolic dysfunction in the lateral portion of the EC of preclinical AD patients. More recently, human alEC volume has been found to relate both to cognitive processing of complex objects^27^ and cognitive decline prior to a clinical diagnosis of amnestic mild cognitive impairment^28^ (aMCI, often considered a prodromal phase towards AD). Finally, in a prior study, we found that older adults were globally impaired at mnemonic discrimination (a likely behavioral readout of pattern separation) of objects compared to young adults, with only very minor differences in spatial discrimination^29^. This is consistent with unique dysfunction of alEC and/or PRC. The phenomenon of age-related object discrimination deficits has also been closely paralleled by recent studies in rats^30,31^. However, key questions remain unresolved. What is the neural basis of object discrimination deficits in older adults? Can alEC dysfunction explain behavioral performance? Given a link between DG/CA3 hyperactivity and object discrimination deficits in prior work^12-14^, is this mechanistically linked to alEC dysfunction?

Here, we addressed these gaps in knowledge by adapting the aforementioned object versus spatial mnemonic discrimination paradigm for use in a high-resolution fMRI experiment. We found two phenomena in the fMRI signal that were associated with object discrimination deficits. The first was hypoactivity of alEC as we hypothesized, and the second was a replication of the previously reported DG/CA3 hyperactivity in older adults^12-14^. Moreover, we find evidence for a shift in the correlational structure between alEC and DG/CA3 activity in aging and suggest that the imbalance across these two regions may partially explain age-related memory deficits.

## Results

### Object discrimination is impaired in older adults

The task is depicted in Figure 1, and a full description can be seen in the *Methods*. Briefly, in initial study phases, participants first studied a series of objects appearing in one of 31 on-screen locations and made an incidental “Indoors or Outdoors?” judgment. They then completed a subsequent test phase for each unique set of objects. Half of the test phases featured exactly repeated objects (object targets) or similar lure objects (object lures) (Fig. 1a), and the other half featured exactly repeated locations (spatial targets) or similar lure locations (spatial lures) (Fig. 1b). Lures of each domain were further divided into high, mid, and low similarity bins. Test judgments for both domains were simply “Same or Different?” with respect to the objects or their locations. Three runs of each type of test were completed. All participants underwent an extensive neuropsychological battery, and all participants included in our analyses were within the norms of their age range (Table 1). Though older adults were outperformed by young participants on several tests, these differences are not uncommon, and no differences were observed on broad measures of cognitive intactness (e.g., the Montreal Cognitive Assessment).

**Table 1:** Demographics and Neuropsychological Tests.

| Measure | Young | Old | Difference (t-test) |
| --- | --- | --- | --- |
| Sample Size | 20 (15F) | 20 (13F) | - |
| Age | 21.75 (4.08) | 73.6 (6.2) | - |
| Montreal Cognitive Assessment | 26.71 (2.59) | 26.14 (2.97) | $p = 0.543$ |
| Craft Story Immediate – Verbatim | 20.89 (5.72) | 18.68 (8.96) | $p = 0.371$ |
| Craft Story Immediate – Paraphrase | 15.06 (2.94) | 14.23 (5.75) | $p = 0.589$ |
| Benson – Immediate | 17 (0) | 16.75 (0.55) | $p = 0.062$ |
| Benson – Delayed | 15.33 (2.5) | 11.2 (2.93) | **$p < 0.001$** |
| Benson - Recognition | 1.89 (3.77) | 0.85 (0.37) | $p = 0.228$ |
| Number Span Forward | 8.94 (2.41) | 8.14 (2.26) | $p = 0.292$ |
| Number Span Backward | 6.72 (1.49) | 7.14 (2.32) | $p = 0.514$ |
| Category Fluency – Animal | 23.72 (5.76) | 22.43 (4.63) | $p = 0.442$ |
| Category Fluency – Vegetable | 13.11 (4.34) | 14.86 (5.09) | $p = 0.261$ |
| Trails A | 17.94 (5.46) | 30.35 (8.52) | **$p < 0.001$** |
| Trails B | 45.39 (9.3) | 78.95 (32.34) | **$p < 0.001$** |
| Craft Story Delayed – Verbatim | 19.22 (5.78) | 14.45 (10.27) | $p = 0.091$ |
| Craft Story Delayed – Paraphrase | 14.78 (3.67) | 12.5 (5.87) | $p = 0.166$ |
| MINT – Correct Uncued | 27 (4.2) | 30.19 (1.89) | **$p = 0.003$** |
| MINT – Correct Semantic Cue | 0.72 (0.96) | 0.29 (0.56) | $p = 0.086$ |
| MINT – Total Correct | 17.21 (14.11) | 22.07 (13.94) | $p = 0.256$ |
| Verbal Fluency – “F” Words | 15 (4.04) | 16.22 (6.16) | $p = 0.562$ |
| Verbal Fluency – “L” Words | 14.72 (6.01) | 15 (5.99) | $p = 0.83$ |
| NAART35 | 16.56 (7.37) | 23.73 (8.3) | **$p = 0.002$** |
| RAVLT – Immediate | 13.72 (1.49) | 10.68 (3.01) | **$p < 0.001$** |
| RAVLT – Delayed | 13.06 (3.6) | 9.73 (3.28) | **$p = 0.001$** |
| RAVLT – Recognition | 13.39 (3.6) | 13.64 (1.84) | $p = 0.719$ |
| WAIS – Vocabulary | 35.22 (9.13) | 39.52 (9.02) | $p = 0.148$ |
| Beck Depression Inventory | 7.89 (6.44) | 5.8 (4.66) | $p = 0.303$ |
| Average Hours Slept (Self-Report) | 7.04 (6.44) | 6.88 (1.43) | $p = 0.744$ |
| Stress Inventory Score | 7.04 (1.35) | 2.8 (1.74) | **$p = 0.021$** |

Data are presented as mean (SD). MINT = Multilingual Naming Test, NAART = North American Adult Reading Test, RAVLT = Rey Auditory Verbal Learning Test, WAIS = Wechsler Adult Intelligence Scale. Group differences were assessed via simple pairwise t-tests. Significant differences are bolded.

**Figure 1:**
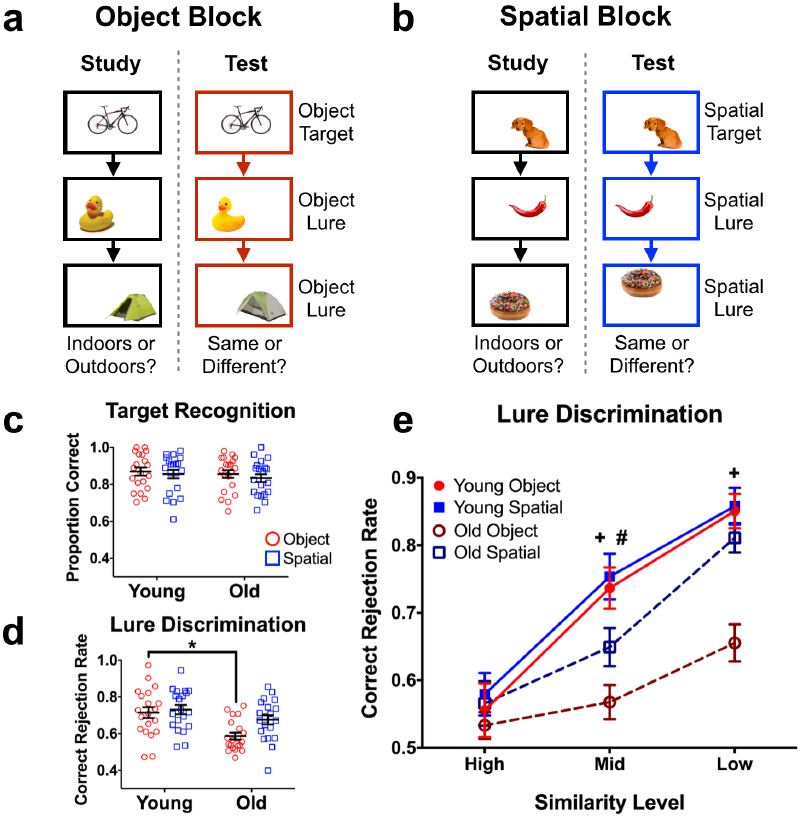
Task schematic and behavioral performance. A) Illustrative diagram of an object block (3 blocks completed). B) Illustrative diagram of a spatial block (3 blocks completed). Objects were smaller relative to screen size in the actual task, and presentation order was randomized across runs (i.e., study and test orders were different) and across participants. Stimuli were presented for 3 seconds, with a 1 second inter-stimulus interval. C) Target hit rates across test domains. No differences were observed. D) Lure discrimination rates across test domains. Older subjects were significantly impaired at object, but not spatial discrimination overall. E) Lure discrimination expanded across similarity bins. Whereas older subjects were relatively impaired at discrimination of mid-similarity spatial lures, they were more globally impaired at object discrimination. Data are shown as mean ± standard error, with each point representing a single subject in panels c and d. (* = Young > Old in panel d; + = Young > Old for object lures in panel E; # = Young > Old for spatial lures in panel E; all *post hoc* tests are reported as p < 0.05 corrected for multiple comparisons.)

Consistent with prior studies, older participants did not differ from the young group on target recognition (F(1,38) = 0.614, p = 0.438). Additionally, there was no difference between recognition of object targets and spatial targets (F(1,38) = 0.614, p = 0.415), and no interaction between age and test domain (object vs. spatial) (F(1,38) = 0.042, p = 0.838). Thus, recognition memory performance was comparable across test domains, and did not differ as a function of age (Fig. 1c). Averaged across similarity levels, lure discrimination was generally poorer in older participants (F(1,38) = 8.62, p = 0.006). We also observed an effect of test domain (F(1,38) = 10.67, p = 0.002) and an interaction between age and test domain for lures (F(1,38) = 5.148, p = 0.029) (Fig. 1d). Šidák-corrected post-hoc comparisons revealed that this difference was largely explained by poorer object discrimination in older adults compared to young (t(76) = 3.649, p_adj_ = 0.001), whereas no group difference was observed for spatial discrimination (t(76) = 1.57, p_adj_ = 0.227).

We next compared performance across similarity levels and between groups, separately examining object and spatial lure discrimination. For object lure discrimination, we found significant effects of similarity (F(2,76) = 72.23, p < 0.001) and age (F(1,38) = 13.36, p < 0.001), and an interaction between similarity and age (F(2,76) = 14.42, p < 0.001) (Fig. 1e). Post-hoc comparisons revealed that older adults were relatively impaired at object lure discrimination at middle (t(114) = 4.167, p_adj_ < 0.001) and low similarity (t(114) = 4.815, p_adj_ < 0.001). For spatial lure discrimination, we observed a significant effect of similarity (F(2,76) = 93.44, p < 0.001) but not age (F(1,38) = 2.425, p = 0.128), and no interaction (F(2,76) = 2.87, p = 0.063) (Fig. 1e). Post-hoc tests revealed that, although there was not a global age effect across spatial lures, older adults were relatively impaired at rejecting middle similarity lures (t(114) = 2.505, p_adj_ = 0.041). In sum, these results demonstrate that cognitively normal older adults were profoundly impaired at mnemonic discrimination of objects with a more subtle impairment in spatial discrimination.

### Age-related hypoactivity in alEC during object discrimination

We subjected all fMRI data to standard preprocessing steps in AFNI, and conducted a general linear model univariate regression analysis with task conditions (including visual perceptual matching trials which served as a baseline task, Supplementary Fig. 1) and nuisance variables (including motion vectors and global signal outside the gray matter) entered as regressors. Given that our hypotheses were specific to MTL regions, we conducted region-of-interest (ROI) analyses in MTL cortex and hippocampal subfields. Briefly, all subjects’ anatomical and functional images were brought into a common template space using a nonlinear registration approach. We then extracted average regression coefficients (beta weights) across anatomical ROIs and compared across task conditions and age groups within regions. ROIs are shown in Figure 2, and processing and analysis of imaging data are detailed in *Methods*.

**Figure 2:**
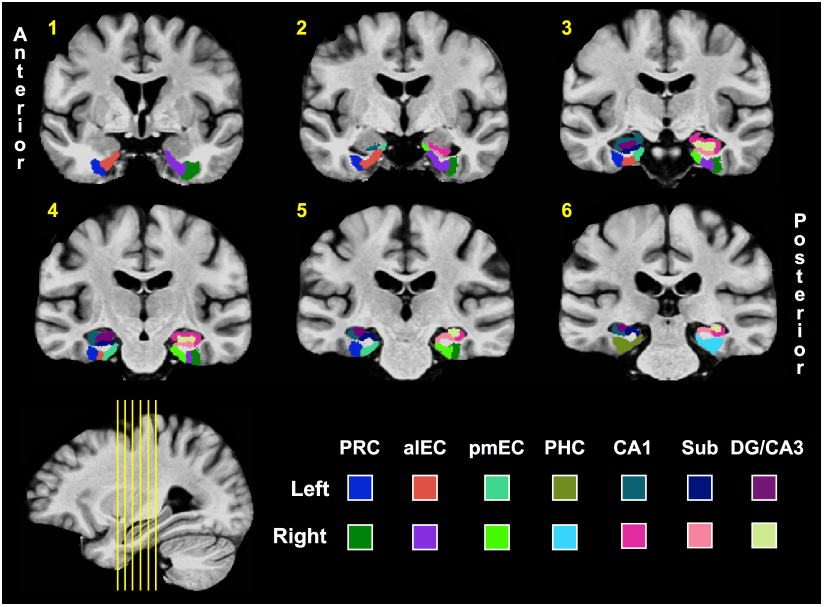
Regions of interest (ROIs). Hippocampal ROIs were based on prior studies^12,13,24,40^, and segmentation of alEC and pmEC was done by hand in accordance with the atlas generated by Maass and colleagues^8^. (PRC = perirhinal cortex; aLEC = anterolateral entorhinal cortex; pMEC = posteromedial entorhinal cortex; PHC = parahippocampal cortex; Sub = Subiculum; DG = dentate gyrus.)

Collapsed across similarity levels, left alEC showed an effect of test domain (F(1,38) = 6.683, p = 0.014) and an interaction between age and test domain (F(1,38) = 5.047, p = 0.031) during lure discrimination. Consistent with our prior work^25^, post hoc comparisons revealed greater engagement of alEC during object discrimination compared to spatial discrimination (t(38) = 3.417, p_adj_ = 0.003) in young participants. A task difference was not observed in older adults, and moreover, left alEC was relatively hypoactive in old compared to young subjects (t(76) = 2.387, p_adj_ = 0.039) (Fig. 3a).

**Figure 3:**
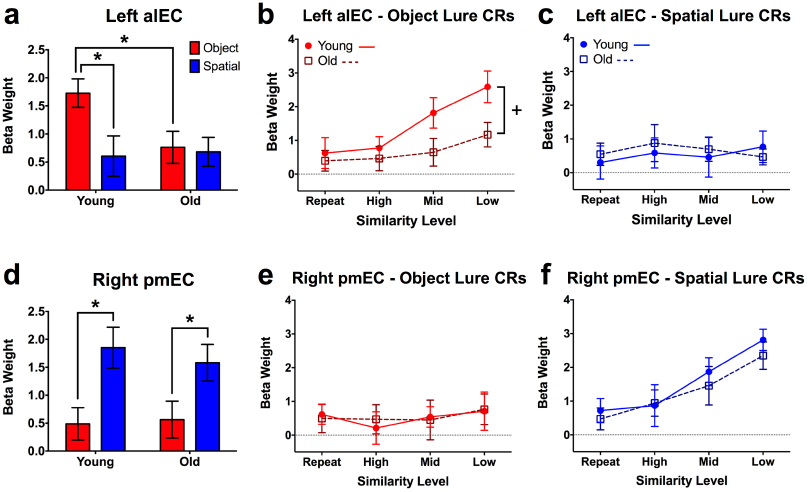
Task and age effects in left alEC and right pmEC. A) Collapsed across similarity levels, older participants show significantly lower activity in left alEC during object discrimination. B) Older adults show significantly lesser gains in alEC engagement with decreasing object similarity compared to young participants. C) alEC shows neither a modulation of spatial lure similarity nor age during spatial discrimination. D) Collapsed across similarity levels, both old and young participants show comparable levels of right pmEC engagement during spatial lure discrimination. E) pmEC shows neither a modulation of object lure similarity nor age during object discrimination. F) For both young and old participants, pmEC is increasingly engaged as lure similarity decreases during spatial discrimination, with no group difference. Data are shown as mean ± standard error (CR = correct rejection; * = Young > Old; + = Significantly different group slopes for the curve of alEC engagement across similarity levels; all post hoc tests are reported as p < 0.05 corrected for multiple comparisons.)

We next analyzed left alEC engagement across trial similarity levels, considering a range from target recognition (exact repetitions, the most similar) to discrimination of high, middle, and low similarity lures (the most dissimilar) (Fig. 3b). During object trials, we observed effects of age (F(1,38) = 4.613, p = 0.038) and similarity (F(3,114) = 7.192, p < 0.001), but no interaction. The data show a marked difference between age groups such that, although left alEC engagement increased as object similarity decreased, the magnitude of this function was considerably blunted in older adults. Post-hoc analyses were conducted using a curve-fitting approach, revealing significantly different slopes across age groups (F(1,156) = 4.136, p = 0.044). In contrast with object trials, the same analysis over spatial trials did not reveal any significant effects (Fig. 3c). Also, critically, age-related functional differences in alEC were independent of differences in cortical volume, as young and old subjects did not differ in this regard (Supplementary Fig. 2) Thus, compared to young participants, older adults showed alEC hypoactivity during object discrimination, which manifested as a blunted response across a range of item similarity independent of any structural differences. Object selectivity was also observed in PRC, though we did not observe age-related hypoactivity there or in any other ROI (Supplementary Fig. 3, Supplementary Tables 1-3).

In right pmEC, the collapsed analyses showed only an effect of test domain (F(1,38) = 15.22, p < 0.001). Post-hoc tests revealed that this was driven by greater task engagement during spatial compared to object discrimination across both young (t(38) = 3.157, p_adj_ = 0.006) and old participants (t(38) = 2.361, p = 0.046) (Fig. 3d). No difference was observed between age groups. Conducting the analyses across similarity levels, no effects were observed in right pmEC during object trials (Fig. 3e). In contrast, during spatial trials, we observed a significant effect of lure similarity (F(3,114) = 12.11, p < 0.001) but neither an effect of age nor an interaction (Fig. 3f). This is consistent with spatial tuning canonically found in rodent studies of the MEC^32^ as well as our prior results in human subjects^25^. A highly similar set of results were observed bilaterally in parahippocampal cortex (PHC; Supplementary Fig. 3, Supplementary Table1-3). Similar to alEC, we did not observe group differences in the volume of pmEC (Supplementary Fig. 2). In sum, whereas alEC was engaged during object but not spatial trials, pmEC was engaged during spatial but not object trials in young adults. Additionally, whereas alEC engagement was blunted in older adults, engagement of pmEC did not differ from young participants during spatial discrimination.

### Age-related hyperactivity in DG/CA3 during lure rejection across domains

We next probed for task and age-related effects in hippocampal subfields. Collapsed across lure similarity in left DG/CA3, we found an effect of age (F(1,38) = 10.89, p = 0.002), but no difference across test domain, and no interaction during lure discrimination (Fig. 4a). Post-hoc comparisons revealed that, unlike the pattern observed in left alEC, engagement of left DG/CA3 was greater for old participants compared to young for both object (t(76) = 2.468, p_adj_ = 0.031) and spatial discrimination (t(76) = 2.78, p_adj_ = 0.014).

**Figure 4:**
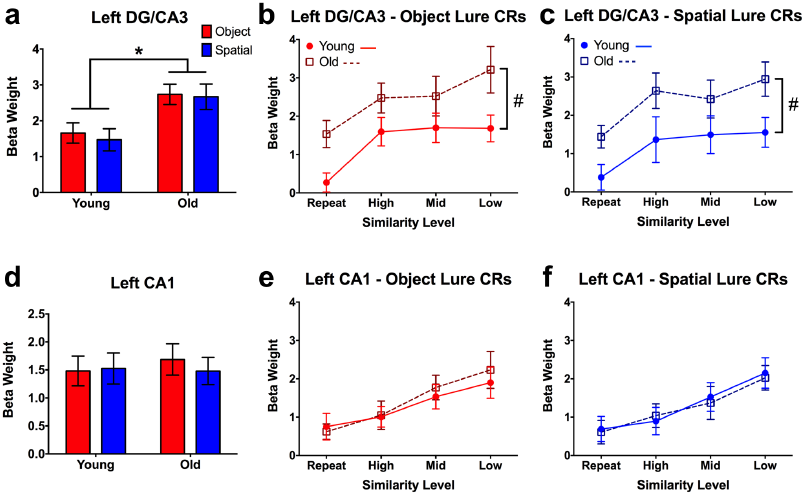
Task and age effects in left DG/CA3 and CA1. A) Collapsed across similarity levels, older participants show significantly higher activity in left DG/CA3 across both test domains. B,C) Compared to young participants, older adults show significantly greater activity in left DG/CA3 across all lure similarity levels during object and spatial discrimination. D,E) Both young and older participants show significantly increasing engagement of left CA1 with decreasing lure similarity, but no group differences. Data are shown as mean ± standard error. (CR = correct rejection; * = Young > Old; # = Significantly different group intercepts (but not slopes) for the curve of DG/CA3 engagement across similarity levels; all *post hoc* tests are reported as p < 0.05 corrected for multiple comparisons.)

Analyzing across similarity bins, we found effects of age (F(1,38) = 10.02, p = 0.003) and lure similarity (F(3,114) = 6.22, p < 0.001) in left DG/CA3 during object trials (Fig. 4b). Similarly, during spatial trials, we found effects of age (F(1,38) = 9.009, p = 0.005) and similarity (F(3,114) = 4.345, p = 0.006) (Fig. 4c). Curve-fitting analysis revealed that while the slopes did not differ between groups, the intercepts differed for both object (F(1,157) = 14.818, p < 0.001) and spatial (F(1,157) = 14.059, p < 0.001) trials. Thus, while the overall “shape” of the functions observed in older adults are similar to the young group, the effect of aging is driven by general hyperactivity in DG/CA3 (across all levels of similarity) across test domains. This is consistent with prior studies in humans^12-14^ as well as rodents^31,33,34^ and monkeys^35^. Similar trends were observed in right DG/CA3, though the effects were not significant (Supplementary Fig. 4, Supplementary Tables 1-3).

We next examined task-related modulations in the CA1 subfield. Though no task or age-related effects were observed when collapsing across similarity levels (Fig. 4d), when considering the full range of lures, we observed significant effects of similarity during object trials (F(3,114) = 8.860, p < 0.001) (Fig. 4e) and spatial trials (F(3,114) = 8.502, p < 0.001) (Fig. 4f). However, in sharp contrast to left DG/CA3, we did not observe effects of age in left CA1. This demonstrates that age-related hyperactivity is not found throughout the hippocampus (see also right CA1 and subiculum, Supplementary Fig. 4, Supplementary Tables 1-3). Moreover, contrasted with the observed linear pattern in CA1, the curvilinear relationship observed in DG/CA3 (driven by a strong response even to highly similar lures) is consistent with a pattern separation transfer function^11^ and is an independent replication of prior work^36^.

### Engagement of DG/CA3 and alEC correlates with behavior

We next asked whether engagement of the key regions discussed here – left DG/CA3 and left alEC – was related to subjects’ discrimination performance. To address this question, we computed simple Pearson correlations between engagement of a given region (beta weights, averaged across similarity levels) and correct rejection rates for object or spatial trials. Correlations were assessed at the level of young subjects, old subjects, and the entire sample collapsed across age. In left DG/CA3, during object lure discrimination, we found a significant negative correlation in older adults (r = −0.454, t(18) = −2.164 p = 0.044) and in the entire sample (r = −0.485, t(38) = −3.423, p = 0.001) (Fig. 5a). Though the relationship was similar in young participants, the correlation was not significant (r = −0.309, t(18) = −1.376, p = 0.186). During spatial discrimination, we found a significant negative correlation between performance and left DG/CA3 across the entire sample (r = −0.37, t(38) = −2.457, p = 0.019), but neither age group featured a significant correlation individually (young: r = −0.283, t(18) = −1.252, p = 0.227) (old: r = − 0.337, t(18) = −1.52, p = 0.146) (Fig. 5b). Thus, in general, greater engagement of DG/CA3 was associated with poorer mnemonic discrimination across test domains.

**Figure 5:**
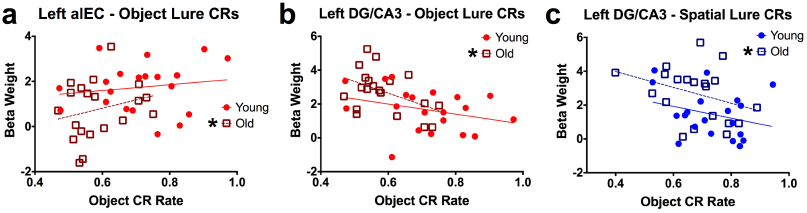
Correlations between left DG/CA3, left alEC, and behavior. A,B) Left DG/CA3 engagement is negatively correlated with object and spatial discrimination performance across participants. Correlations tended to be stronger in older adults. C) Left alEC engagement is positively correlated with object discrimination performance, a relationship driven largely by older participants. (CR = correct rejection; * beside group icon = significant correlation.)

We next examined brain-behavior relationships in the left alEC. During object discrimination, engagement of left alEC was positively associated with behavioral performance in older adults (r = 0.46, t(18) = 2.195, p = 0.042) and across the entire sample (r = 0.41, t(38) = 2.774, p = 0.009) (Fig. 5c). The correlation was not significant in young adults (r = 0.154, t(18) = 0.661, p = 0.517). Thus, unlike the negative relationship observed in DG/CA3, greater engagement of alEC was associated with better object discrimination performance.

### Relationship between DG/CA3 and alEC is disrupted in aging

The above regional results suggest that DG/CA3 hyperactivity and alEC hypoactivity may be related conditions that are associated with object discrimination deficits. To examine whether these conditions are related, we asked whether alEC and DG/CA3 may be coupled in young and older adults during object discrimination trials. Rather than simply correlate activity between the regions, we computed a normed ratio of DG/CA3 engagement relative to alEC engagement (sum of DG/CA3 and alEC divided by difference of DG/CA3 and alEC, with greater values being associated with a bias toward DG/CA3 activity). Young subjects had a mean ratio near zero, suggesting balanced engagement of DG/CA3 and alEC. Conversely, the ratio was significantly higher for older adults (t(18) = 3.139, p = 0.006), indicating greater DG/CA3 activity relative to alEC activity (Fig. 6a). Moreover, we found a significant negative correlation between ratio scores and object discrimination performance in older adults (r = −0.51, t(18) = −2.533, p = 0.021). Though the relationship was qualitatively similar in young subjects, there was not a significant correlation (r = −0.213, t(18) = −0.927, p = 0.366). In sum, in addition to disrupting the engagement of alEC and DG/CA3 individually, aging seems to imbalance the relationship between the regions during mnemonic discrimination of objects in a way that predicts impaired performance.

**Figure 6:**
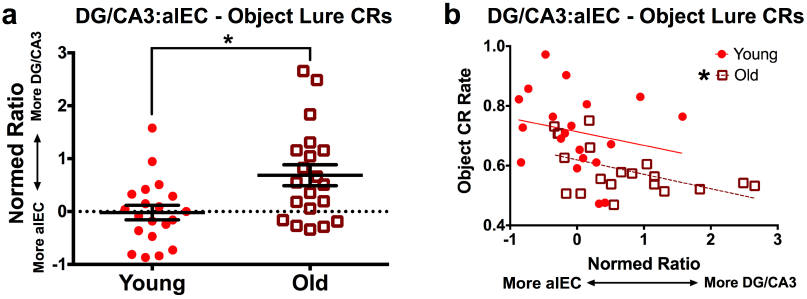
Normed ratios of left DG/CA3 vs. alEC activity across groups and relationship with behavior. A) Older participants have a significantly higher ratio value than young adults, indicating greater DG/CA3 activity relative to alEC. Conversely, young participants have a near-zero mean ratio, indicating relatively balanced engagement of both regions. B) Object discrimination performance negatively correlates with this normed ratio in older adults, indicating greater DG/CA3 activity relative to alEC is associated with poorer object mnemonic discrimination. Despite a qualitatively similar relationship in young participants, the relationship does not yield a significant correlation. (CR = correct rejection; * in panel A = significant group difference; * beside group icon in panel B = significant correlation).

## Discussion

In this study, we observed severe age-related impairments in mnemonic discrimination of objects, contrasted with subtler impairments in spatial discrimination (Fig. 1). This was coincident with specific hypoactivity in left alEC (Fig. 3) and hyperactivity in left DG/CA3 (Fig. 4). Moreover, activity in alEC was positively correlated with object discrimination performance, whereas activity in DG/CA3 was negatively correlated with mnemonic discrimination across object and spatial trials (Fig. 5). Finally, a normed ratio of DG/CA3 and alEC activity demonstrated a skew toward DG/CA3 engagement at the expense of alEC in older adults, which correlated with the extent of object discrimination deficits (Fig. 6). Importantly, these older participants were asymptomatic in terms of dementia or major indices of cognitive decline.

The domain-selective functional dissociation we observed between alEC and pmEC is consistent with our prior findings^25^ as well as resting state analyses^22,23^. Our results also accord with recent reports of alEC structural properties relating to object perception and cognitive outcomes^27^. However, we build significantly on these lines of research by providing, for the first time to our knowledge, evidence for age-related dysfunction of alEC *in vivo*. It is important to consider that these participants were cognitively normal. Additionally, our multi-domain approach allowed us to dissociate subtly impaired spatial discrimination from more drastically impaired object discrimination. This implies that even relatively “normal” aging might feature functional decline of alEC alongside selectively disrupted mnemonic discrimination of objects. Importantly, it has been widely hypothesized that functional disruption of MTL circuits precedes structural degradation by a considerable amount of time^37-39^. Indeed, typical age-related memory decline is not in itself associated with significant loss of cells in the MTL^40^. Thus, with this paradigm, we may be uncovering subtle changes to the MTL that may confer vulnerability on the aging brain.

Pathological hippocampal hyperactivity is now fairly well-established in age-related cognitive decline across rodents^31,33,34^, nonhuman primates^35^, and humans^12-14^. This hyperactivity is relatively specific to the CA3 subfield of the hippocampus, and loss of GABAergic inhibitory interneurons may be a contributor to this phenomenon^35^. A recent set of studies across transgenic mice^41^, rats^34,42^, and humans^43^ with age-related cognitive impairment related hippocampal hyperactivity to memory deficits. Reduction of hyperactivity with a low-dose of levetiracetam (LEV), an anti-epileptic drug targeting excitatory neurotransmission, largely rescued memory deficits as well as CA3 hyperactivity (combined DG/CA3 in human subjects), and a more recent study in humans found that EC hypoactivity was also normalized by the treatment^44^. Our results are therefore well in line with these findings, and extend them to suggest a specific role for alEC (as opposed to the entire EC) as well as the potential importance of the relationship between alEC and DG/CA3.

This raises a key question: how do hypoactivity in alEC and hyperactivity in DG/CA3 relate? It must of course be noted that fMRI is unable to weigh in on the dynamics of neuronal circuits or the directionality of shared signaling. However, a recent study in rodents by Maurer et al.^31^ offers some insight. In addition to projections to DG, perforant path input from the lateral EC projects directly to CA3. In their experiment, Maurer and colleagues found that old rats with object discrimination deficits featured CA3 hyperactivity as well as a greater overall proportion of active CA3 cells, which was directly influenced by inputs from lateral EC. As CA3 is more excitable than DG even in healthy young animals, an overall drop in alEC signaling could potentially lead to an imbalance in the EC-hippocampal circuit such that the CA3 autoassociative networks is reinforced. Thus, in the case of objects, this could bias the hippocampus away from novel encoding of incoming sensory information (i.e., through pattern separation) and more toward generalization based on stored representations (i.e., through pattern completion)^11^. As the signal from EC is corrupted on its way into the hippocampus, and as the signal passes hierarchically through the network, the mismatch between EC and the hippocampal representation may become amplified during CA3 autoassociation. Another possibility, which is not mutually exclusive, is that hyperactivity in CA3 could impose a stressful condition that leads to retraction of pyramidal cell apical dendrites and retrograde degeneration of the perforant path from alEC. Perforant path loss has been demonstrated reliably in aged rodents^45-47^, as well as more recently in humans^13,14^ and occurs in the absence of cell loss in the region. This would suggest that CA3 hyperactivity may be a driving factor in alEC hypoactivity, which would be consistent with the impact of LEV administration which is assumed to target hyperactivity, but additionally rescues alEC hypoactivity perhaps by re-tuning hippocampal dynamics and allowing the CA3 to process alEC input which in turn strengthens alEC signaling.

Our data and those of Maurer and colleagues^31^ suggest that dysfunction to the alEC-CA3 circuit may be specifically linked to this age-related mnemonic discrimnination deficits observed. The exact mechanisms and temporal order of these events remains an important question for future work to continue to address. However, the observed disruption of alEC and DG/CA3 dynamics, namely the ratio of engagement between regions, offers a unique and potentially powerful avenue for understanding healthy versus pathological aging^38^. For instance, one might hypothesize that individuals with AD-related pathology (such as amyloid and tau accumulation) or those along the road to major cognitive decline will feature ratios more skewed toward DG/CA3 activity relative to healthy controls, or that this skew will emerge earlier. Thus, the ratio measure may serve as a potential biomarker for clinical interventions. Future studies will be needed to examine these links, and longitudinal experiments tracking participants over time can evaluate the predictive power of such a candidate biomarker.

In conclusion, we provide a mechanistic basis for object discrimination impairments in aging – an imbalance in the alEC-DG/CA3 circuit characterized by reduced signaling in the alEC coupled with increased signaling in DG/CA3. This work has important implications for future assessments of MTL regions, as well as specific mnemonic functions as we characterize different aging trajectories.

## Methods

### Subjects

Twenty-six young adults and 25 older adults were initially recruited for the study from the UC Irvine and greater Orange County community. One young participant and one older participant were excluded from analysis due to confusion with the task. An additional two young adults and one older participant were excluded due to excess motion in the scanner. Finally, three young participants and three older participants were excluded due to poor signal in the medial temporal lobes (particularly the entorhinal cortex – see *Image Preprocessing* below). This yielded a final sample of 20 young adults (15 female, range = 18-31 years, mean = 21.75 years, SD = 4.08 years) and 20 older adults (13 female, range = 64-89 years, mean = 73.6 years, SD = 6.2 years). All subjects were screened for neurological conditions (e.g., history of stroke or mental illness). Subjects gave written informed consent in accordance with the UC Irvine Institutional Review Board, and were compensated for their participation.

A neuropsychological battery was administered to characterize our sample (Table 1). Importantly, broad assays of cognitive impairment (e.g., the Montreal Cognitive Assessment) did not differ as a function of age. We did observe group differences in the Trails A and B tests (sensitive to executive function and task flexibility), the Benson Delayed Recall test (sensitive to memory for complex objects, though the Recognition test was unaffected), the RAVLT Immediate and Delayed Recall tests (sensitive to verbal memory, though the Recognition test was unaffected), the NAART35 (sensitive to verbal intellectual ability), and MINT Uncued (sensitive to verbal fluency). However, declining performance on these particular tests are not uncommon in typical aging, and even our lowest performing subjects were not significantly outside their age-matched range.

### Behavioral Task

The task was an optimized version of a prior paradigm^29^ which we optimized for the scanner. Participants completed six blocks of study and test, with three blocks testing memory for object identity and the other two testing memory for spatial locations Fig. 1). To facilitate ease of the task for older adults, subjects completed three blocks of either the object or spatial task consecutively before switching to the other. Though a randomized block delivery would be ideal, pilot testing suggested that some older participants struggled with switching between tasks flexibly. Importantly, the order of task completion was counterbalanced across our sample.

Stimuli consisted of colored images of common objects appearing on a 7 × 5 spacing scheme (accommodating a widescreen display). Objects were displayed for 3 seconds, with a 1 second inter-stimulus interval. Study and test sequences each consisted of 50 trials. Of these 50 trials, 42 consisted of images of objects. Both study and test runs included 8 trials of a visual perceptual matching task (Supplementary Fig. 1), which were used as a baseline for task-driven activity estimates (see *MRI Data Analysis* below). Briefly, during these conditions, subjects saw two blurred dots and were asked whether they were equally dark (the actual probability of equal shading was 50%, and the differences in opacity when present varied from 5%-25%). During object test blocks, targets were identical to studied objects whereas lures were objects that were perceptually similar, but not identical (Fig. 1a). Similarity bins for object lures were based on *a priori* similarity indices validated by prior studies^15,36^. Specifically, we used bins 2, 3, and 4 (out of the full range of 1-5) reported by these studies. During spatial test blocks, targets were studied objects occupying the same grid space whereas lures were studied objects occupying a different location than the original location (Fig. 1b). Similarity bins for spatial lures were based on prior work^25,29,48^, matched to object similarity bins resulting in 2, 3, and 4 grid moves for high, mid, and low spatial similarity respectively. Unlike several of our prior studies, we did not include completely novel items at test.

Subjects were aware that their memory was to be tested following study blocks, but specific non-mnemonic judgments during study to foster attention to stimuli. Critically, judgments did not differ as a function of task. That is, all study blocks featured “Indoors or Outdoors?” judgments, and all test blocks featured “Same or Different?” judgments (referring to the object itself in object blocks, and the location in spatial blocks). Responses were recorded via button-press. Objects were unique to each block, and each space on the grid (excluding corners) was equally likely to be occupied. Within a block, trial order was completely random. The task was programmed in Python (version 2.7) using PsychoPy^49,50^.

### Image Acquisition

Neuroimaging data were acquired on a 3.0 Tesla Philips Achieva scanner, using a 32-channel sensitivity encoding (SENSE) coil at the Neuroscience Imaging Center at the University of California, Irvine. A high-resolution 3D magnetization-prepared rapid gradient echo (MP-RAGE) structural scan (0.75 mm isotropic voxels) was acquired at the beginning of each session: repetition time (TR)=11 ms, echo time (TE)=5.03ms, 231 slices, 0.65mm isotropic, field of view (FOV)=231.174×240×150.150. Functional MRI scans consisted of a T2*-weighted echo planar imaging (EPI) sequence using blood-oxygenation-level-dependent (BOLD) contrast: TR=2500 ms, TE=26 ms, flip angle=70 degrees, 39 slices, 84 dynamics per run, 1.8 x 1.8 mm in plane resolution, 1.6 mm slice thickness with a 0.2 mm gap, FOV=180×77.8×180. Slices were acquired as a partial axial volume and without offset or angulation (Supplementary Fig. 5). Four initial “dummy scans” were acquired to ensure T1 signal stabilization. A total of 12 functional runs were acquired for each participant, 6 study phases and 6 test phases (3 each of study and test for object and spatial blocks), except for one older participant who had only two usable runs of the spatial memory test (however, a sufficient number of trials were acquired from the first two runs). Each functional run lasted 4 minutes and 30 seconds. For each subject, T2-weighted scans and ultrahigh-resolution diffusion-weighted scans were also acquired, though they are not analyzed here.

### Image Preprocessing

All neuroimaging data were preprocessed and analyzed using Analysis of Functional NeuroImages (AFNI, version 17.2.00)^51^ on GNU/Linux and Mac OSX platforms. Analyses largely took place in accordance with the standardized afni_proc.py pipeline. EPIs were corrected for motion (3dvolreg) and slice timing (3dTshift), masked to exclude voxels outside the brain (3dautomask), and were smoothed (3dmerge) using a 2.0mm Gaussian FWHM kernel. Motion correction parameters were saved into text files for later use in linear regression (see *MRI Data Analysis* below). Each run was also despiked to further reduce the influence of motion on the data (3dDespike). Functional scans were aligned to each subject’s skull-stripped MP-RAGE (align_epi_anat.py). We used Advanced Normalization Tools (ANTs)^52^ to warp each individual participant's MP-RAGE structural scan into our custom in-house high-resolution 0.75mm isotropic template using SyN nonlinear registration. Parameters from these warps were used to also warp functional scans into template space for group ROI analyses (see *ROI Segmentation* and *MRI Data Analysis* below). Masks were resampled to match the resolution of the fMRI data (2.0mm isotropic) and were further masked to exclude partially sampled voxels within and across runs (3dcalc).

Prior to further analysis, we took two steps to ensure that data quality – particularly in the EC, given signal dropout issues in the region – was sufficient for each participant. First, we calculated the ratio of EPI voxels in the alEC and pmEC ROIs to overall voxels in the ROI, which served as an index of how much dropout occurred. Subjects missing greater than 25% of alEC or pmEC voxels were excluded. Second, we examined the signal-to-noise ratio of the time series (TSNR) for each subject. Overall TSNR was adequate, but several participants featured considerably lower values in the MTL. Examples of “good” and “poor” MTL tSNR can be seen for young and older participants in Supplementary Figures 6 and 7, respectively. We excluded subjects whose TSNR in the alEC or pmEC was >50 for greater than 25% of voxels in the ROI. These steps ultimately resulted in 3 young and 3 older participants being excluded from further analysis.

### ROI Segmentation

We defined ROIs in the medial temporal lobes (MTL) based on our established protocol1^2,25^. Briefly, segmentation of hippocampal subfields was conducted in accordance with the SY protocol reported in Yushkevich and colleagues^53^ though the CA1-subiculum boundary was updated to reflect recent efforts at harmonizing across hippocampal segmentation protocols^54^. As we did not have hypotheses regarding the longitudinal axis of the hippocampus, we did not split hippocampal ROIs into anterior and posterior segments. The most notable addition to the protocol was segmentation of alEC versus pmEC. Whereas we previously segmented the EC into lateral and medial portions^25^, here we incorporated the segmentation of Maass et al.^22^ on the basis of their results and related findings of Navarro-Schroder et al.^23^. Briefly, these two groups independently found resting time-series correlations between PRC and PHC to respective alEC and pmEC subregions, and Maass and colleagues^22^ generated a freely-available alEC and pmEC segmentation on their group template. We resampled their data to the resolution of ours, and hand-segmented the ROIs into our template slice-by-slice in the coronal plane using the Maass et al.^22^ template as a reference. All ROIs are displayed in a series of 6 slices in Figure 2. Our group template and ROI mask are available upon request.

### MRI Data Analysis

Only test data are included in the analyses here. We constructed a general linear model (GLM) with regressors for all trial types: target hits, target misses, lure rejections, lure false alarms, and perceptual task trials. We additionally included regressors for six motion vectors derived from the motion correction preprocessing step (x, y, z, pitch, roll, yaw) as well as non-response trials. There were too few target misses to analyze activity during those trials, and give that our hypotheses were fairly specific to lure rejections, we also did not analyze activity during lure false alarms. These steps were conducted in AFNI using 3dDeconvolve. Deconvolution of the hemodynamic response was done using tent functions covering stimulus onset to 15 seconds after onset with 6 estimator functions distributed across this time window. Motion parameters were entered into the model as explicit regressors to reduce the influence of motion on task-related parameter estimates, and non-response trials were entered to exclude these ambiguous trials from serving as a baseline condition. Additionally, vectors modeling temporal drift were entered as regressors covering first and second-order polynomials. We explicitly modeled perceptual matching trials (Supplementary Fig. 1) as the baseline for general linear tests of our task conditions, as this serves as a fairly robust “zero” point for comparing conditions in MTL regions^55^. For all functional runs, TRs with motion exceeding 0.5mm frame displacement (but below our exclusion threshold of 3mm) were censored from analyses, as well as the immediately preceding and following TRs. Finally, global signal from the ventricles and white matter was excluded from gray matter voxels using ANATICOR^56^. These fairly rigorous “data scrubbing” procedures were employed to attempt to exclude the potential effects of head motion on activation profiles^57^.

Final beta weights entered into second-level analyses consisted of the average of the first three estimator functions (targeted to capture the peak of the BOLD response). These beta weights were extracted from ROIs (3dmaskave), and ANOVAs were used to probe significant effects of age, test domain, and lure similarity across regions (see *Statistical Testing and Data Availability* below). Though we employed an exclusively ROI-based approach given our *a priori* hypotheses about the data, we note that, in general, image contrasts of the effects reported here (e.g., greater alEC activity in young compared to older participants) are significant at the level of a group t-test (Supplementary Fig. 8). Moreover, the robustness of these effects can be seen by examining lure discrimination vs. target hit contrasts at the level of individual subjects’ beta weight maps (Supplementary Fig. 9).

The normed ratio between left DG/CA3 and alEC (Fig. 6) was calculated as the sum of DG/CA3 and alEC beta weights divided by the difference of DG/CA3 and alEC beta weights. This approach has been used in prior studies^58^ and yields a measure of the relative engagement between two regions. In the present formulation, alEC beta weights was added to or subtracted from those of DG/CA3. Consequently, a greater value indicates a ratio more heavily in favor of DG/CA3 engagement, whereas a smaller value indicates a ratio more heavily in favor of alEC engagement.

### Cortical Volume Estimates for alEC and pmEC

Each participant’s MTL was segmented using a multi-atlas label fusion (MALF) approach^59^. This approach uses multiple segmentation atlases, which consisted of twenty expert-labeled example brains (hippocampal labels were applied for use in prior publications, and alEC/pmEC labels were added by Z.M.R.). During the MALF procedure, all segmentation atlases are registered to a target image (i.e., each participant’s brain). The MALF weighting scheme for choosing the ROI label at each voxel in the target image simultaneously maximizes atlas similarity within a local neighborhood of the target image while minimizing informational redundancy amongst the atlas set. The consequence of this is that for any given ROI, the atlas whose brain structure most closely matches the participant is weighed most heavily in labeling that ROI in a subject’s space, which accounts for a great deal of structural variability across subjects. Cortical volume measures in alEC and pmEC were extracted as the overall volume (in mm^3^) across the ROI as labeled on the subject’s brain, which were aggregated across subjects. Results are plotted in Supplementary Figure 2.

### Statistical Testing and Data Availability

Image preprocessing and analyses were conducted in AFNI (version 16.0) using standard functionality (noted throughout the *Methods* section), and custom shell scripts were created to batch routines. These scripts as well as the structural and functional MRI data are available upon request. Statistical tests and figures were performed with GraphPad Prism (Version 6.07), and the relevant data file is available upon request. All statistical tests were two-tailed.

For both behavior and fMRI data, analyses collapsed across similarity bins were carried out using two-way mixed ANOVAs, with age as a between-subjects factor and test domain as a within-subjects factor. For analyses across similarity bins for a given domain, tests were carried out using two-way mixed ANOVAs, with age as a between-subjects factor and similarity as a within-subjects factor. As we performed ROI-based analyses and did not conduct any statistical comparisons *across regions*, assessing effects within one region should be statistically independent of testing within another region^60^. We therefore did not adjust our global significance threshold for detecting main effects in these ANOVAs. Pairwise post hoc comparisons were conducted via the Šidák correction^61^. Briefly, this sets a familywise error rate of 0.05 by accounting for the number of comparisons. Relationships between behavior and fMRI measures were assessed with Pearson correlations. Contrasts between the slopes and intercepts of correlations across age groups were conducted using curve fitting analyses, implemented as simple linear fits in Prism.

## Acknowledgements

We thank our participants for volunteering to take part in our research. We thank Matt Tsai, Vishali Kapoor, and Diana Salama for assistance with data collection. We also thank Rebecca Stevenson, Maria Montchal, Corey Fernandez, and Craig Stark for helpful discussions. This work was supported by US National Institute on Aging P50 AG05146 and NIA R21 AG049220 (to M.A.Y.), as well as support for Z.M.R. provided by US National Science Foundation Graduate Research Fellowship DGE-1232825, NIA Training Grant T32 AG-000096, and fellowships from the UCI Chancellor’s Club and the ARCS/Roche Foundations.

## Author Contributions

Z.M.R. and M.A.Y. designed the research. Z.M.R. and J.A.N. collected data. J.A.N. and E.A.M. recruited participants. Z.M.R., N.J.T., and D.D. conducted analyses. Z.M.R. and M.A.Y. wrote the paper.

